# Prescribed versus wildfire impacts on exotic plants and soil microbes in California grasslands

**DOI:** 10.1101/2021.09.22.461426

**Authors:** Sydney I Glassman, James WJ Randolph, Sameer S Saroa, Joia K Capocchi, Kendra E Walters, M. Fabiola Pulido-Chavez, Loralee Larios

## Abstract

1. Prescribed fire is often used as a management tool to decrease exotic plant cover and increase native plant cover in grasslands. These changes may also be mediated by fire impacts on soil microbial communities, which drive plant productivity and function. Yet, the ecological effects of prescribed burns compared to wildfires on either plant or soil microbial composition remain unclear.
2. Here, we investigated the impacts of a spring prescribed fire versus a fall wildfire on plant cover and community composition and bacterial and fungal richness, abundance, and composition in a California grassland. We used qPCR of 16S and 18S to assess impacts on bacterial and fungal abundance and Illumina MiSeq of 16S and ITS2 to assess impacts on bacterial and fungal richness and composition.
3. Wildfire had stronger impacts than prescribed fire on microbial communities and both fires had similar impacts on plants with both prescribed and wildfire reducing exotic plant cover but neither reducing exotic plant richness. Fungal richness declined after the wildfire but not prescribed fire, but bacterial richness was unaffected by either. Yet increasing char levels in both fire types resulted in reduced bacterial and fungal biomass, and both fire types slightly altered bacterial and fungal composition.
4. Exotic and native plant diversity differentially affected soil microbial diversity, with native plant diversity leading to increased arbuscular mycorrhizal fungal richness while exotic plant diversity better predicted bacterial richness. However, the remainder of the soil microbial communities were more related to aspects of soil chemistry including cation exchange capacity, organic matter, pH and phosphorous.
5. *Synthesis and applications*. Understanding the different ecological effects of prescribed fires and wildfires on plant and soil communities are key to enhancing a prevalent management action and to guide potential management opportunities post wildfires. Our coupled plant and soil community sampling allowed us to capture the sensitivity of the fungal community to fire and highlights the importance of potentially incorporating management actions such as soil or fungal amendments to promote this critical community that mediates native plant performance.

## Introduction

Land managers increasingly need to grapple with the effects of fire on their management objectives, particularly in the Western United States, where prescribed burns and wildfires are becoming more frequent (Running 2006; Westerling *et al*. 2006; Ryan, Knapp & Varner 2013). Comparisons between the effects of prescribed burns and wildfires have centered on forests where long-term fire suppression has altered the baseline conditions and can create undesirable negative effects (Ryan, Knapp & Varner 2013). However, similarities between prescribed burns and wildfires in other systems, such as grasslands, have yet to be fully explored. Within grasslands, leveraging insights from cultural burning practices, land managers are using prescribed burns as a management tool to reduce the negative impacts of exotic species and increase native plant establishment and performance (Blackburn & Anderson 1993; McKemey *et al*. 2020). Prescribed fires typically occur in spring, under higher moisture conditions that minimize fire spread and intensity (Ryan, Knapp & Varner 2013). In contrast, wildfires in the America West historically occur during the summer and fall, under drier fuel conditions (Stephens & Collins 2004), thus leading to more intense burns. This difference in burn intensity may be a key factor in the beneficial management outcomes of prescribed burns (Knapp, Estes & Skinner 2009), but native plant species performance is not consistently enhanced in prescribed burns and can be negatively affected by wildfires (Alba *et al*. 2015). Therefore, more studies are needed that compare the ecological effects of prescribed burns and wildfires in grasslands to improve the use of fire as a management tool.

The impacts of fire for land management have classically focused on native plant recovery but have overlooked the impacts that the soil microbial communities may have on vegetation recovery. Soil microbes are critical drivers of all major biogeochemical cycles (Crowther *et al*. 2019) and plant diversity and productivity (van der Heijden *et al*. 1998). Thus, their resilience, or rate of return to the undisturbed state (Shade *et al*. 2012), after fire may be critical to the regeneration of aboveground vegetation. Fungi, in particular, likely play critical roles in plant regeneration since up to to 90% of plant species are associated with mycorrhizal fungal symbionts that increase access to nutrients in exchange for carbon (Brundrett & Tedersoo 2018). Arbuscular mycorrhizal fungi (AMF) are particularly important in driving grassland dynamics (van der Heijden *et al*. 2006), and AMF often co-occur with soil bacteria and fungi that may be critical to plant growth, plant productivity, and the cycling of nutrients (Garbaye 1994; Yuan *et al*. 2021). Despite the importance of plant-AMF coupling in mediating plant successional dynamics (Kardol, Bezemer & van der Putten 2006; Cheeke *et al*. 2019), the impacts of fires on soil microbes in grasslands remains understudied (Pressler, Moore & Cotrufo 2019).

How prescribed burns versus wildfires affect soil microbial resilience may critically impact native plant regeneration. Fires typically reduce soil microbial biomass (Dooley & Treseder 2012) and richness (Pressler, Moore & Cotrufo 2019), and alter the richness and composition of mycorrhizal fungal symbionts (Dove & Hart 2017). A meta-analysis of belowground impacts of fires found that only 12% of studies focused on grasslands (Pressler, Moore & Cotrufo 2019), even though roughly 80% of fires globally occur in grasslands each year (Leys *et al*. 2018). While a review found that prescribed fires and wildfires have similar impacts on fungal biomass in grasslands (Pressler, Moore & Cotrufo 2019), the impacts of prescribed fires versus wildfires on grassland bacterial biomass, microbial richness, or composition remain unknown. The lower fuel load in grassland typically results in fast-moving low severity wildfires. Soil microbes are more resilient to less severe fires (Holden *et al*. 2016), thus, fires in grasslands, whether prescribed or wildfire, might not be severe enough to significantly alter soil microbial communities.

Exotic plant species, which are often the impetus for prescribed burns (Knapp, Estes & Skinner 2009), may create soil legacies that limit the recovery of native species (Kulmatiski *et al*. 2008). These soil legacies may interact with disturbances such as fire to synergistically reduce native plant performance (Suding *et al*. 2013). For example, many exotic plant species succeed and become invasive through reduced reliance on mycorrhizal symbionts (Pringle *et al*. 2009) or by suppressing native symbiotic communities in an invaded area (Mummey & Rillig 2006; Vogelsang & Bever 2009; Zubek *et al*. 2016), thus promoting the growth of invasive plant species over resident native species. Alternatively, exotic plant species may alter the soil microbial community by enhancing pathogens (Eppinga *et al*. 2006) or promoting soil communities that enhance resource acquisition (Hawkes *et al*. 2006). These dynamics can result in a positive feedback loop (Callaway *et al*. 2004; Kulmatiski *et al*. 2008) and may interact with disturbances to further promote the performance of exotic species (Suding *et al*. 2013). They can also limit the outcomes of restoration efforts for native plant species (Lankau *et al*. 2014). Therefore, understanding the resilience of the soil microbial community in an invaded grassland is key to identifying and refining further management efforts.

California grasslands are an ideal system to explore these dynamics. Prescribed fires are an important management tool for reducing the dominance of exotic species in once biodiverse native perennial grasslands (Menke 1992; Dyer 2002; DiTomaso *et al*. 2006). When used as a restoration tool, prescribed burns have had mixed effects in helping to eliminate noxious weeds, especially in systems where invasive annual grasses within the genera *Avena* and *Bromus* outcompete native perennial bunchgrass, thus failing to promote native plant species establishment (Holmes & Rice 1996; Meyer & Schiffman 1999; D’Antonio *et al*. 2002; Corbin & D’Antonio 2010). These annual grasses can differentially impact biogeochemical cycling and soil communities for their benefit (Hawkes *et al*. 2005; Eviner & Hawkes 2008; Vogelsang & Bever 2009). However, soil amendments to offset the impacts of these invaders can result in positive native plant responses (Sandel, Corbin & Krupa 2011; Emam 2016). Together, these dynamics suggest that a closer investigation of how soil microbial communities are changing with fire may provide some insight into improving the efficacy of grassland management.

Here, we explore the impacts of a prescribed fire versus a wildfire that burned through remnant perennial grassland in Southern California. Prescribed fires, which typically occur in spring, are used as a management tool to reduce nonnative species cover and to retain cover of the native perennial bunchgrass *Stipa pulchra* in Southern California grasslands (Valliere *et al*. 2019), but summer and fall wildfires also typically occur in California grasslands every 2-6 years (Fryer & Luensmann 2012). We coupled time-series sampling of the soil microbial community with plant recovery data to ask how prescribed fire versus wildfire impacts: 1) native versus exotic plant cover and richness 2) soil bacterial and fungal biomass and richness 3) soil bacterial and fungal composition over time, and finally, 4) how does AMF and soil microbial regeneration relate to native plant regeneration?

## Materials and Methods

### Description of field site

Field work was conducted at the Santa Rosa Plateau Ecological Reserve, Riverside County, CA, USA in the location of the Burro Burn unit and the Tenaja wildfire (**Table S1)**. The reserve consists of expanses of grassland intermixed with chaparral, coastal sage scrub, and oak woodlands (Valliere *et al*. 2019) situated in a Mediterranean climate with dry summers and where most precipitation falls from October through May. Precipitation for the 2018-19 growing season was approximately 648 mm and 506mm for 2019-2020. Both soils are Alfisols with the soil at the Tenaja site belonging to the Vallecitos series and the soilat the Burro site in the Murrieta series (https://casoilresource.lawr.ucdavis.edu/gmap/). The Burro prescribed fire took place on May 23, 2018 and burned 0.6 km^2^. The Tenaja wildfire took place from Sept 4, 2019 through Sept 14, 2019 and burned 7.8 km^2^. Both sites were located in grasslands dominated by invasive annual grasses from the genus *Bromus*.

### Sampling Methods

Three replicated transects were established, with plots 5m outside the burn line (A), and inside the burn line 5m (B), 20m (C), 100m (D) and 200m (E) (**Fig. 1)**. Transects were established based on proximity to contiguous habitat opposite from the burn section outlined by a firebreak. All lines were set up to be as far as possible from the access road, to cover as much area of the burn unit as possible, and to represent varying distances from the burn edge that might be of relevance to plants and microbes dispersing in from unburned edges. The design was initially established to test the effects of time since fire and distance from burned edge on recovery of bacterial, fungal and plant communities post-fire to pre-fire. However, we observed no effect of distance from burned edge and dropped that variable from the analyses presented here. For the Tenaja fire, we added an additional 2 unburned plots in between the three transects for a total of 5 unburned plots and 12 burned plots. Therefore, we had 15 plots within the Burro prescribed fire burn unit and 17 within the Tenaja wildfire burn scar.

**Figure 1.**
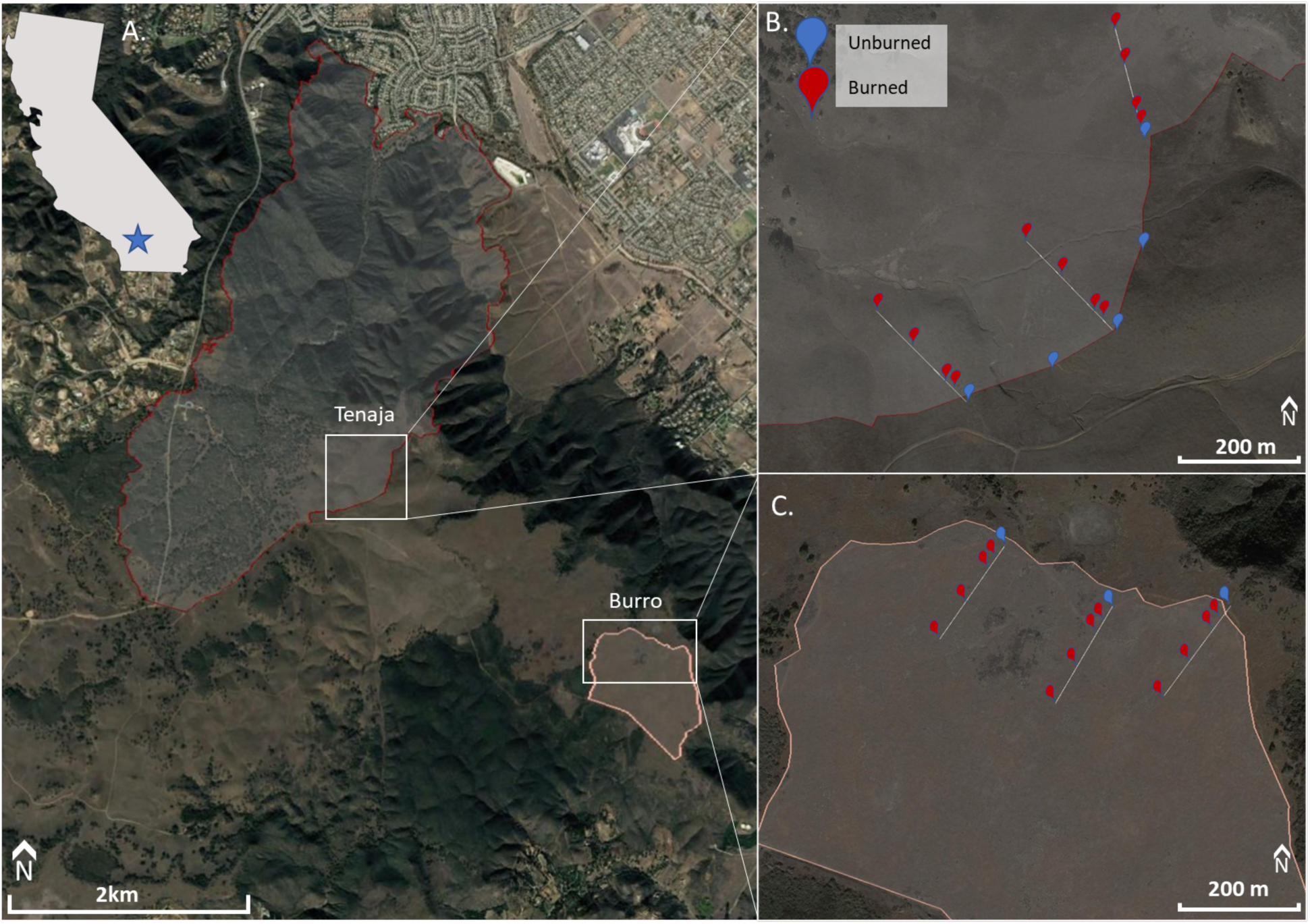
Map of study site with both fire perimeters displayed (A) and insets show the sampling transects within the Tenaja wildfire (B) and Burro prescribed burn (C). Samples were collected 5m outside the burn (blue), and 5m, 20m, 100m, and 200m (red) within the burn. Two additional unburned samples were sampled between transects within the Tenaja wildfire.

These plots were sampled at three time points post-fire. Within the prescribed fire, soils were collected on 6-2-2018 at 2 weeks post-fire (1^st^ post-fire), on 11-28-2018 at 6 months post-fire (Nov), and on 5-6-2019 at 12 months post-fire (May). The wildfire soils were sampled on 11-5-2019 at 2 months post-fire (1^st^ post fire), on 11-19-2019 at 2.5 months post-fire (Nov), and on 5-13-2020 at 7 months post-fire (May). The first time point captured the immediate effects of fire on the soil microbial community. Unfortunately, due to access constraints for the wildfire, this time frame differs between both burn units. The remaining two sampling points paralleled growing season phenology to account for the effects of intra-seasonal variation in precipitation on soil communities.

At every time point, for each plot, three ∼5cm deep subsamples (8cm diameter and 10 cm deep releasable bulb planter filled half depth) were collected haphazardly from the 1m^2^ plot and combined for a single homogenized sample per plot. Between plots gloves and soil corers were sterilized with 70% ethanol. All soils were stored on ice in the field and stored in -80°C freezer that night. A total of 45 soil samples were collected for the prescribed fire (3 transects x 5 plots x 3 time points) and 51 soil samples were collected for the wildfire (3 transects x 5 plots + 2 additional unburned plots x 3 time points). At the first post-fire sampling, the amount of bare ground, rock, litter, char material and live plant cover was estimated for each 1m^2^ plot. Litter was identified as any plant growth from previous growing seasons that had not decomposed, and char material was identified as any ash or blacked charred residue present on the ground surface post fire. We used initial char as a proxy for fire severity as these post-fire residues may mediate nutrient pulses associated with fire and can be linked to fire severity (Neary *et al*. 1999). In spring of the first post-fire year, the plant community composition was visually estimated in each plot. The cover of each species was estimated such that percent cover could be over 100 to capture species cover in the overlapping layers. Observers used reference squares of known percentages to help estimate cover and increase consistency across different observers.

### Soil Chemistry

Soil from 32 soil samples from the first post-fire sampling were selected for nutrient analysis, which consisted of 15 samples from 2 weeks post-fire for prescribed fire and 17 samples at 2 months post-fire for the wildfire as this provided the best temporal comparison between the two sites (i.e. the closest sampling after the fire). Soils were air-dried in a fume hood, and sent to A&L Western Laboratories, Inc. (Modesto, CA, USA) for analysis (http://www.al-labs-west.com/fee-schedule.php?section=Soil%20Analysis; soil test suite S1B) including ppm of sulfur (S), potassium (K), magnesium (Mg), calcium (Ca), and sodium (Na), soil pH, cation exchange capacity (CEC), hydrogen concentration, organic matter (OM) in lbs/acre, and phosphorous (weak Bray and Sodium Bicarbonate-P; ppm). To minimize redundancy in some of the soil variables, we dropped variables that were estimates of similar pools (e.g. weak Bray and sodium bicarbonate -P) and variables with strong correlations that indicated similar dynamics (e.g. both Mg and Ca were strongly correlated with CEC at r>0.8, therefore just CEC was kept). This reduced our soil variables to OM, phosphorous (Sodium Bicarbonate-P; ppm), soil pH, K, Na, CEC, and S. These seven variables were analyzed in a principal components analysis to get two axes that described the soil attributes of the sites. The first axis, which captured 30.74% of the variation, described plots with high soil pH, P, and K availability at negative values of PC1 but high Na values at positive values of PC1. The second axis, which captured 26.96% of variation, described plots across a CEC, S, and OM gradient where negative values indicated plots with high CEC and positive values plots with low CEC (**Fig. S1**).

### DNA extractions, PCRs, and Illumina MiSeq sequencing

To identify microbial biomass, richness, and composition, DNA was extracted from 0.25g soil using Qiagen DNeasy PowerSoil Kits following the manufacturer’s protocol and stored at -20°C for subsequent analysis. Extracted DNA was amplified using the primer pair ITS4-fun and 5.8s (Taylor *et al*. 2016) to amplify the ITS2 region of the Internal Transcribed Spacer (ITS), which is the internationally recognized barcode for fungi (Schoch *et al*. 2012), and the primer pair 515F-806R to amplify the V4 region of the 16S rRNA gene for archaea and bacteria (Caporaso *et al*. 2012) using the Dual-Index Sequencing Strategy (DIP) (Kozich *et al*. 2013). Although the 515F-806R amplifies both archaea and bacteria, from here on we say only bacteria for simplicity since bacterial reads were so dominant (**Fig. S2)**. We conducted polymerase chain reaction (PCR) in two steps, each with 25µl aliquots. The first PCR amplified gene-specific primers, and the second PCR ligated the DIP barcodes for Illumina sequencing with detailed methods in **Methods S1**. We sequenced with Illumina MiSeq 2×300bp at the University of California-Riverside Institute for Integrative Genome Biology. Sequences were submitted to the National Center for Biotechnology Information Sequence Read Archive under BioProject PRJNA761493.

### Bacterial and fungal biomass

For all soil samples, bacterial and fungal biomass as a proxy of abundance were estimated by qPCR copy number. Bacterial biomass was estimated based on bacterial 16S rRNA genes using the Eub338/Eub518 primer set (Fierer *et al*. 2005) and fungal biomass was estimated based on fungal 18S rRNA genes using the FungiQuant-F and FungiQuant-R primer set (Liu *et al*. 2012). See **Methods S2** for methodological details.

### Bioinformatics

All Illumina MiSeq data were processed using Version 2019.10 of QIIME2 pipeline (Bolyen *et al*. 2019). Forward and reverse adaptors were removed with the cutadapt trim-pair function, and reads were denoised, trimmed to remove low quality regions, and forward and reverse reads were merged using the dada2 denoise-paired functions. Bacterial reads were then tested against the Silva 132 16S classifier (Quast *et al*. 2013) and fungal reads against the UNITE classifier (Köljalg *et al*. 2005). For fungi, all reads not identified to Kingdom Fungi were removed, and for bacteria, all mitochondrial and chloroplast genes were removed.

### Statistical Analysis

All statistical analyses were conducted in R version 4.0.2 (R Core Team 2017). To evaluate the plant community response to the prescribed burn (Burro) and wildfire (Tenaja), we ran mixed effects model for species richness, Shannon’s diversity, and cover of the exotic and native plant community as a function of burned (burned, unburned) and fire (prescribed, wildfire). Sampling plot was nested within transect as a random effect. To evaluate microbial community responses to the two different fires, we ran mixed effect models on bacterial and fungal richness and abundance (as estimated by copy number) separately for each fire as a function of initial char, or the char measured at the first time-point following fire, time and their interaction. Since we assume that initial char measurements are most representative of fire severity for the plants and microbes, we analyze all effects as a function of initial char cover. Bacterial and fungal richness were estimated as observed species number after rarefying all samples to same sequencing depth (21,921 for bacteria, 6,338 for fungi) to account for uneven sequencing depth. We included sampling plot as a random effect and specified an autoregressive correlation structure to account for the lack of independence of samples over time. We compared differences among levels by least square means using the “emmeans” package in R (Lenth *et al*. 2020). Bacterial and fungal abundance were natural log transformed to meet assumptions of normality for data analysis.

To assess recovery of the microbial communities to the fires, we first calculated a Bray-Curtis dissimilarity matrix for the bacterial and fungal communities after rarefaction. For each fire, we used this distance matrix as the response variable in a PERMANOVA (permutational multivariate analysis of variance) with fire (burned/unburned), time, and their interaction as predictor variables. We visualized these differences using a nonmetric multidimensional scaling (NMDS) plot. We also compared the dispersion of the microbial communities between burned and unburned sites for each fire. These analyses were done using the “vegan” package in R (Oksanen *et al*. 2012).

We took a model fitting approach to relate soil properties and plant community attributes to bacterial and fungal communities. We began with a global model that included initial char estimates, soil PCA1 and PCA2, and exotic plant diversity and cover as fixed effects, and transect as a random effect. For the plant community attributes, we selected exotic plant diversity and cover after adjusting for correlations among various community attributes such as richness, cover, and diversity of the total community, native species and exotic species. These models were therefore run with just the May timepoint data for bacterial, fungal, and plant communities. We conducted stepwise model selection and selected the model with the lowest AIC. We also repeated this analysis with AMF richness as the response variable. The model fitting was done using the “MuMIn” package (Barton 2020). Given that the AMF community may be more sensitive to native plants, we conducted an additional analysis where we ran a mixed effects model for AMF richness as a native plant diversity and fire (prescribed, wildfire). Sampling plot was nested within transect as a random effect.

## Results

### Summary description of fire severity

The burned plots within the prescribed burn site were covered on average by 44% char and still had on average 35% cover of intact litter. The burned plots within the wildfire site had on average 74% char cover and no litter was present. In contrast, litter in unburned plots made up about 78% of the plot cover in the first post fire sampling in the prescribed burn site and about 77% cover in the wildfire site. By the spring peak growing season sampling, char was no longer observed in the prescribed burn burned plots and was on average 10% in the wildfire burned plots.

### Summary description of site plants

We observed 65 unique plant species within the prescribed burn site and 41 within the wildfire site with an average species richness of 13 species/m^2^ within the prescribed and 8 species/m^2^ at the wildfire site. The most dominant species in the wildfire site regardless of fire treatment were exotic annual grasses, mainly *Bromus diandrus* and *Avena fatua;* these species made up on average 64% cover in the unburned plots and 49% in burned plots. In the prescribed burn, there was a greater number of dominant species. In the unburned plots, the dominant species were exotic annual grasses (*B. hordeacous, Festuca myuros, F. perennis*) and the native forb, *Calandrinia menziesii;* these species ranged in cover from 13% to 23% cover. In the prescribed fire burned plots, the dominant species each ranged in cover from 9% to 18% cover and consisted of the exotic grass (*F. myuros*), exotic forbs (*Erodium botrys* and *E. cicutarium*), and native forbs (*Calandrinia menziesii, Deinandra fasciculata). Stipa pulchra*, the native perennial bunchgrass that is a focal species for management at the Preserve, averaged ∼1% cover at both sites and was absent in the prescribed fire unburned plots.

### Summary description of site soil bacteria and fungi

Overall, we identified 6,338 fungal and 21,921 bacterial Amplicon Sequence Variants (ASVs), which are similar to operational taxonomic units (OTUs), and approximate microbial species (Glassman & Martiny 2018). After rarefaction, there were 6,602 bacterial and 1,991 fungal ASVs in the prescribed burn and 3,884 bacterial and 1,995 fungal ASVs in the wildfire site at the first time point post-fire, but 6,705 bacterial and 1,948 fungal ASVs were observed in the prescribed burn and 3,699 bacterial and 1,439 fungal ASVs were observed in the wildfire site in the May following the fire. Mean bacterial and fungal richness and abundance were both higher in prescribed burn than wildfire (**Table S2)**. The prescribed burn and wildfire plots were both dominated by the bacterial phyla Proteobacteria and Actinobacteria, followed by Acidobacteria, Bacteroidetes, Verrucomicrobia, Chloroflexi, Planctomycetes, Gemmatimonadetes, and Cyanobacteria (**Fig. S3**). Fungi were always dominated by the phylum Ascomycota regardless of site, followed by Basidiomycota, then Glomeromycota (**Fig. S4)**, with larger turnover from unburned to burned at the family level (**Fig. S5)**.

### Effect of prescribed versus wildfire on exotic and native plant species richness, diversity, and cover

Exotic plant species richness and Shannon diversity were greater in the prescribed than wildfire site (richness: F_1,27.3_= 7.79, p<0.01, **Fig. 2A**; div: F_1,16.6_= 17.05, p<0.001;) and did not differ between the burned and unburned plots. Both the prescribed fire and wildfires reduced exotic species cover, which was greater in the unburned plots (F_1,28_ = 4.15, p=0.05), but did not vary between the two fire sites (F_1,28_ = 0.60, p=0.44, **Fig. 2B**). Native species richness, diversity, and cover were all greater in the prescribed fire site compared to wildfire site and were not affected by burning (**Table S3**; **Fig. 2**).

**Figure 2.**
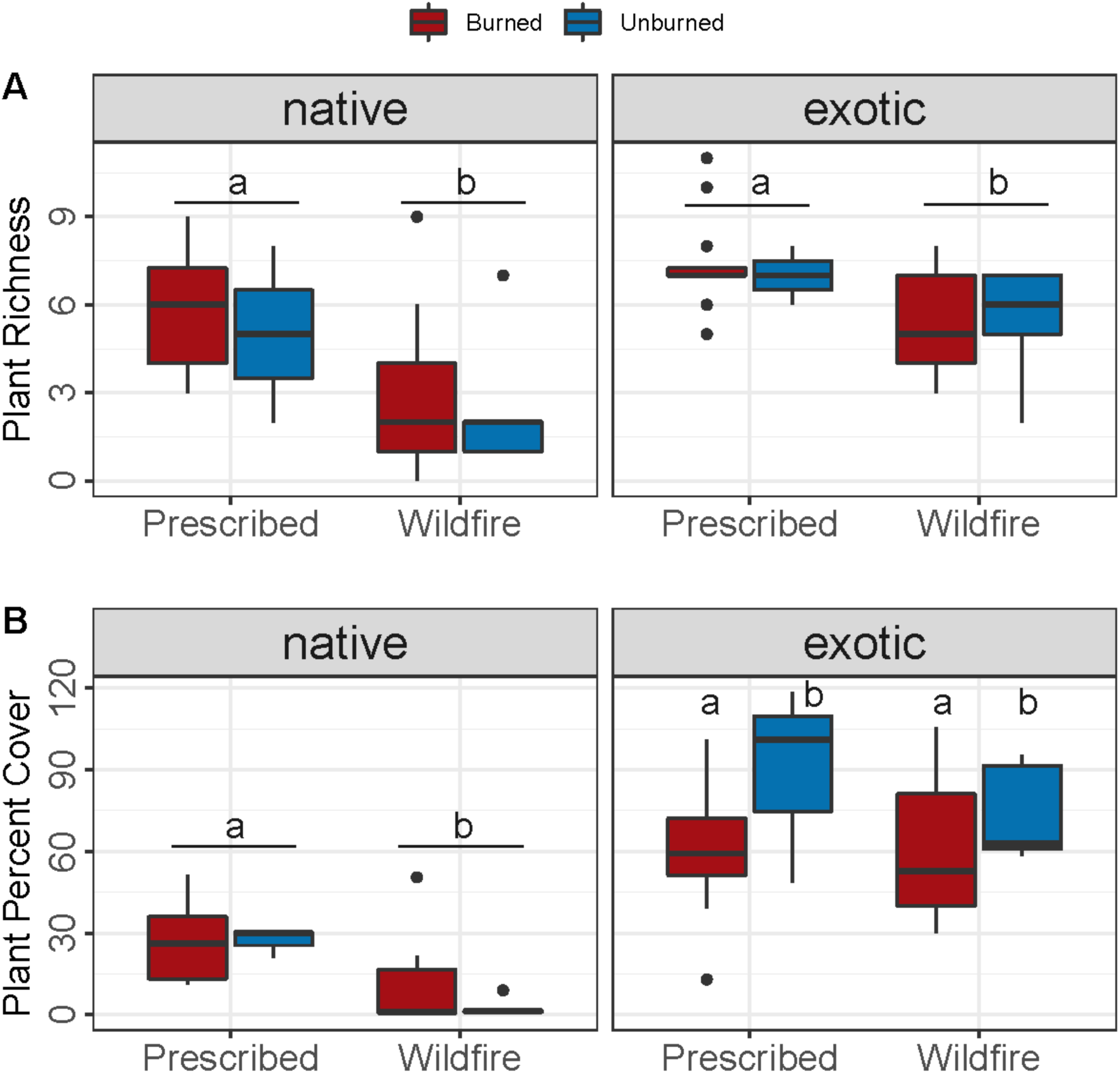
Plant community metrics of species richness (A) and percent cover (B) for native and exotic species in burned and unburned plots within the Burro prescribed burn and Tenaja wildfire sites. Segments and letters indicate significant differences between Burro and Tenaja at p<0.05. Pairwise letters denote the significant effect of burning on exotic species cover at p<0.05.

### Effect of prescribed versus wildfire on soil bacterial and fungal richness

Within the prescribed fire, bacterial richness was not influenced by the initial char but did vary over the three sampling points (Char F_1,13_=0.39, p=0.54; Time F_2,26_=3.5, p<0.05; **Fig. 3A**), where bacterial richness decreased significantly between Nov and May but did not vary among the other time points (post hoc Nov-May p< 0.05). Within the wildfire, bacterial richness was similarly not affected by char but did vary over time, with richness being the greatest in Nov and not differing between the 1^st^ post-fire sampling and May (Char F_1,29_=0.17, p=0.69; Time F_2,29_=7.3, p<0.01; post hoc 1^st^-Nov p<0.05; Nov-May p<0.01; **Fig. 3B**). Fungal richness was not impacted by the prescribed fire such that it did not vary with initial char or over time (Char F_1,13_=0.15, p=0.7; Time F_2,26_=0.11, p=0.89; **Fig. 3C**). Within the wildfire, fungal richness decreased with increasing initial char (Char F_1,29_=22.4, p<0.001) and gradually decreased over time (F_2,29_=16.4, p<0.001, **Fig. 3D**).

**Figure 3.**
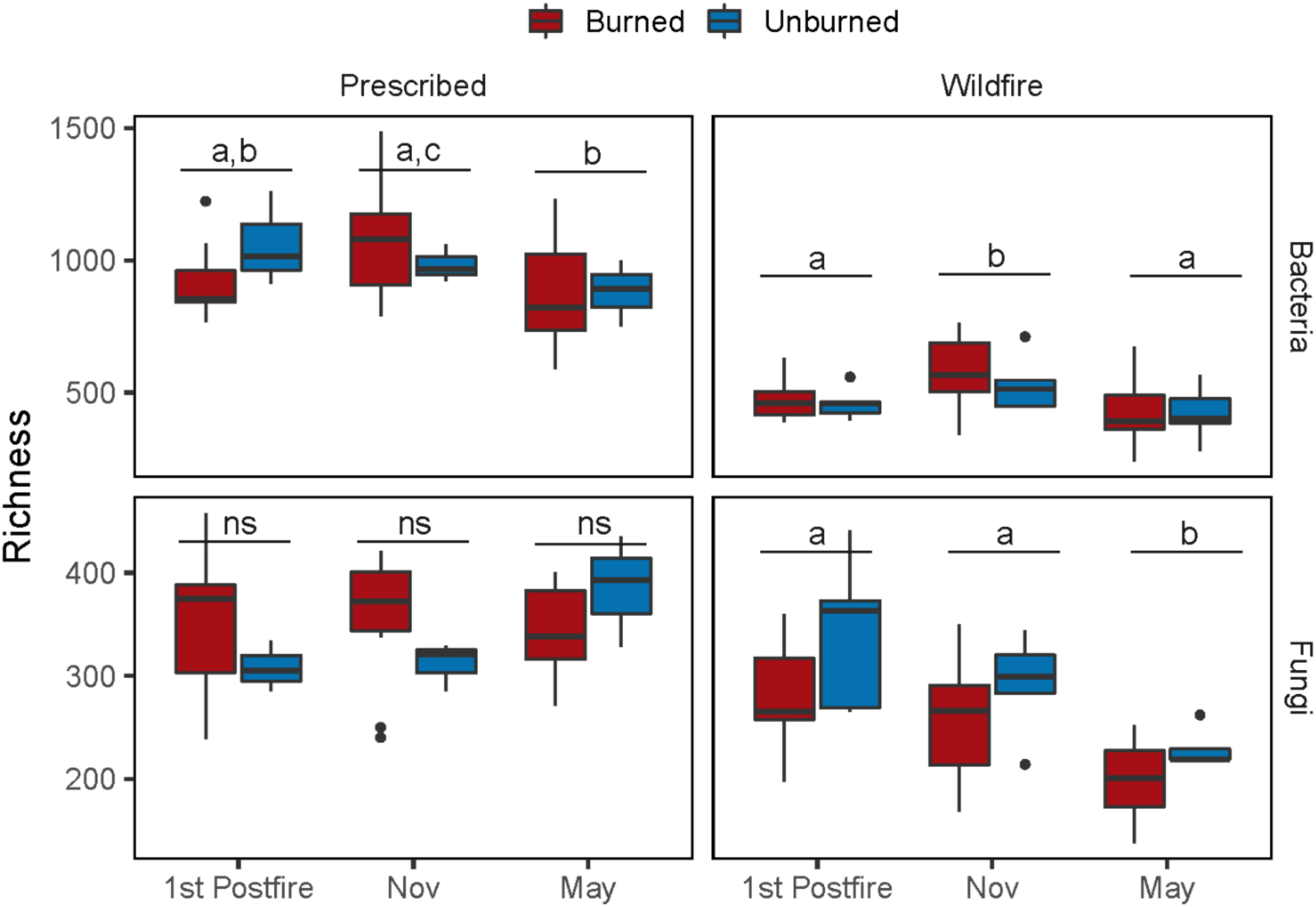
Bacterial and fungal richness for both Burro prescribed fire and Tenaja wildfire over time in burned and unburned plots. Segments and letters indicate post-hoc differences for the main effect of time at p<0.05.

### Effect of prescribed versus wildfire on soil bacterial and fungal abundance

Bacterial and fungal abundance were more responsive than richness to initial char across both fires (**Table S4**). In the prescribed fire, the relationship between bacterial abundance and initial char tended to shift depending on the sampling point (Char x Time F_2,26_=2.86, p=0.08; **Fig. 4A**). For instance, there was no relationship between abundance and char at the first post-fire sampling, but it shifted to positive in November (i.e. abundance increased with char) and then negative in spring. Conversely in the wildfire, bacterial richness slightly decreased with increasing char (Char F_2,29_=7.8, p<0.01, **Fig. 4B**). Bacterial abundance did increase in the wildfire over the sampling periods (Time F_2,29_=29.1, p<0.0001; post hoc at p<0.05 1^st^ Post fire=Nov<May). Fungal abundance also varied with initial char depending on the sampling point within the prescribed burn (Char x Time F_2,26_=5.88, p<0.01; **Fig. 4C**). For the first sampling post-fire, fungal abundance increased with increasing char but for the subsequent two sampling periods the relationship was negative. Within the wildfire, fungal abundance showed a similar trend to bacterial abundance with a small decrease with increasing char (Char F_2,29_=6.3, p<0.02, **Fig. 4D**). While the relationship between char and fungal abundance did not change with time, average fungal abundance tended to decrease from the initial post-fire sampling point to May (Time F_2,29_=6.04, p<0.01; post hoc at p<0.05 1^st^ Post fire=Nov<May).

**Figure 4.**
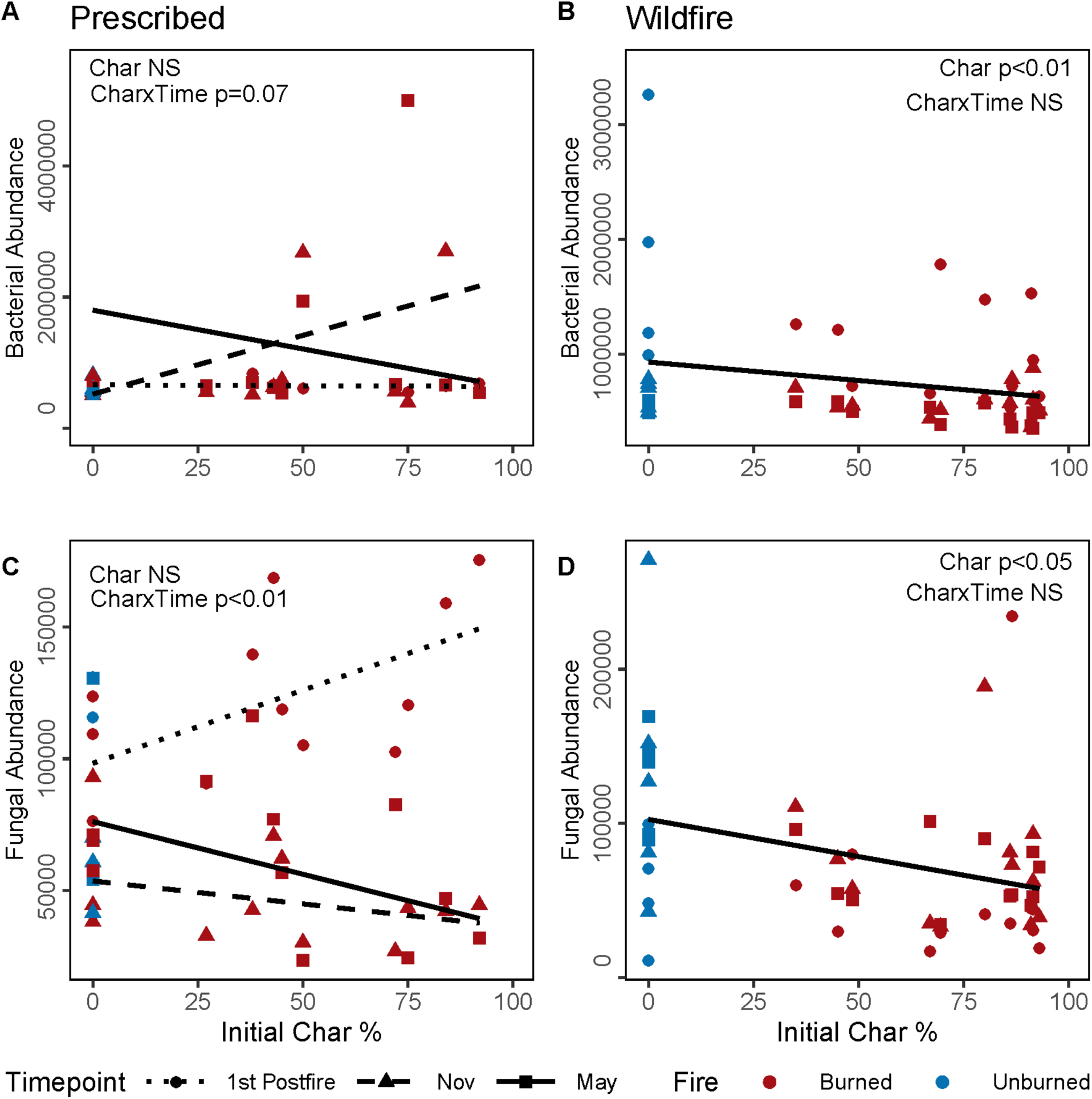
Bacterial abundance, estimated as 16S copy number, at the A) Burro prescribed fire and the B) Tenaja wildfire. Fungal abundance, estimated as 18S copy number, at the C) Burro prescribed fire and the D) Tenaja wildfire. Lines represent significant relationships for either main effect of char on abundance or two-way interactions of Char and Time.

### Effect of prescribed versus wildfire on soil bacterial and fungal composition

Bacterial composition differed to the same degree between burned and unburned plots in both the prescribed burn (PERMANOVA: F_1,39_= 1.45, p<0.05, R^2^=0.03; **Fig. 5A)** and wildfire (F_1,45_= 1.46, p<0.05, R^2^=0.03; **Fig. 5B)** but not over time (PERMANOVA: prescribed-Time F_2,39_= 1.23,p=0.07, R^2^=0.06; wildfire-Time F_2,45_= 1.07, p=0.26, R^2^=0.04). Fungal composition varied between burned and unburned plots both in the prescribed burn (PERMANOVA: F_1,39_= 1.82, p<0.01, R^2^=0.04; **Fig. 5B)** and wildfire (F_1,45_= 2.39,p<0.01, R^2^=0.05; **Fig. 5C**), with slightly more compositional change in the wildfire than prescribed fire. Similar to bacteria, fungal composition did not vary over the three sampling periods in either fire (PERMANOVA: prescribed: F_2,39_= 1.06, p=0.34, R^2^=0.05; wildfire: F_2,45_= 1.05,p=0.35, R^2^=0.04). The variation in community composition differed between the burned and unburned plots for fungal communities in both fires (prescribed, p<0.05, wildfire, p< 0.001) but not for bacterial communities (prescribed, p=0.27, wildfire, p=0.70).

**Figure 5.**
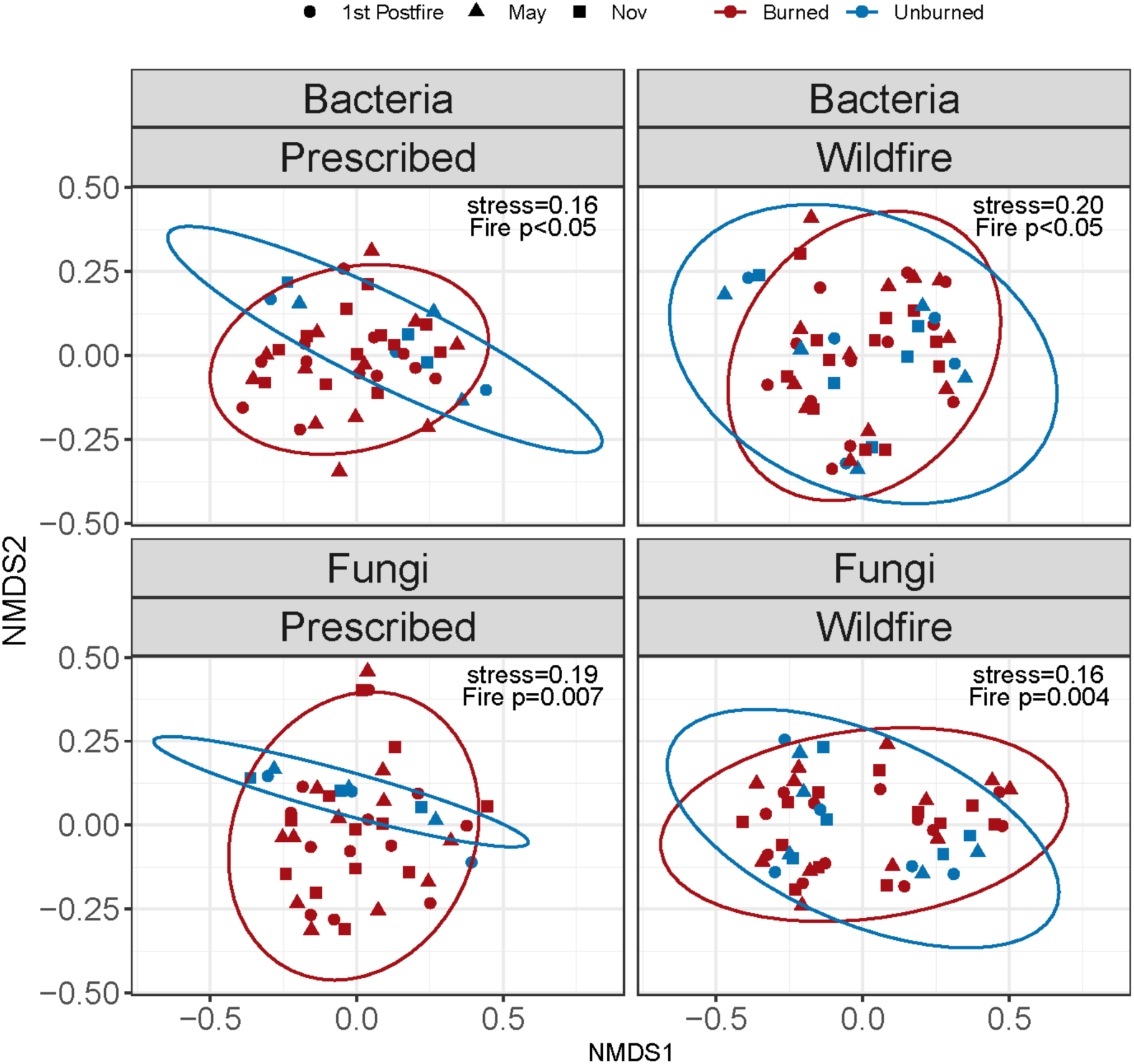
Nonmetric Multidimensional Scaling ordination of burned versus unburned soil microbial composition for bacteria and fungi in the Burro prescribed fire versus Tenaja wildfire at the three time points.

### Predictors of bacterial and fungal richness in Spring

Bacterial and fungal richness were predicted by different soil properties at each site (**Fig. 6**). Within the prescribed burn, bacterial and fungal richness increased with decreasing cation exchange capacity (CEC) and organic matter (OM) (**Fig. 6 A, B**, blue dots). Bacterial richness was also influenced by the plant community such that it increased with increasing exotic plant richness. For the wildfire, bacterial and fungal richness both responded to PCA1 but in different directions. Bacterial richness decreased as soil pH and phosphorus concentrations decreased (**Fig. 6A**, red dots); however fungal richness increased as soil pH and phosphorus concentrations decreased (**Fig. 6B**, red dots). AMF richness on its own was not related to soil properties or to the exotic plant community in either fire (**Fig 6C**). AMF richness followed similar patterns to native plant richness in that the prescribed site (31<u>±</u>5) had higher AMF richness than the wildfire site (17.3<u>±</u>4.3; Fire, F_1,26_=9.76, p<0.01). While AMF richness was not related to exotic plant diversity, it was marginally related to native plant diversity within the wildfire, such that AMF richness increased with increased native plant diversity (Fire x Native F_1,27_=3.14, p=0.09; **Fig. 7)**.

**Figure 6.**
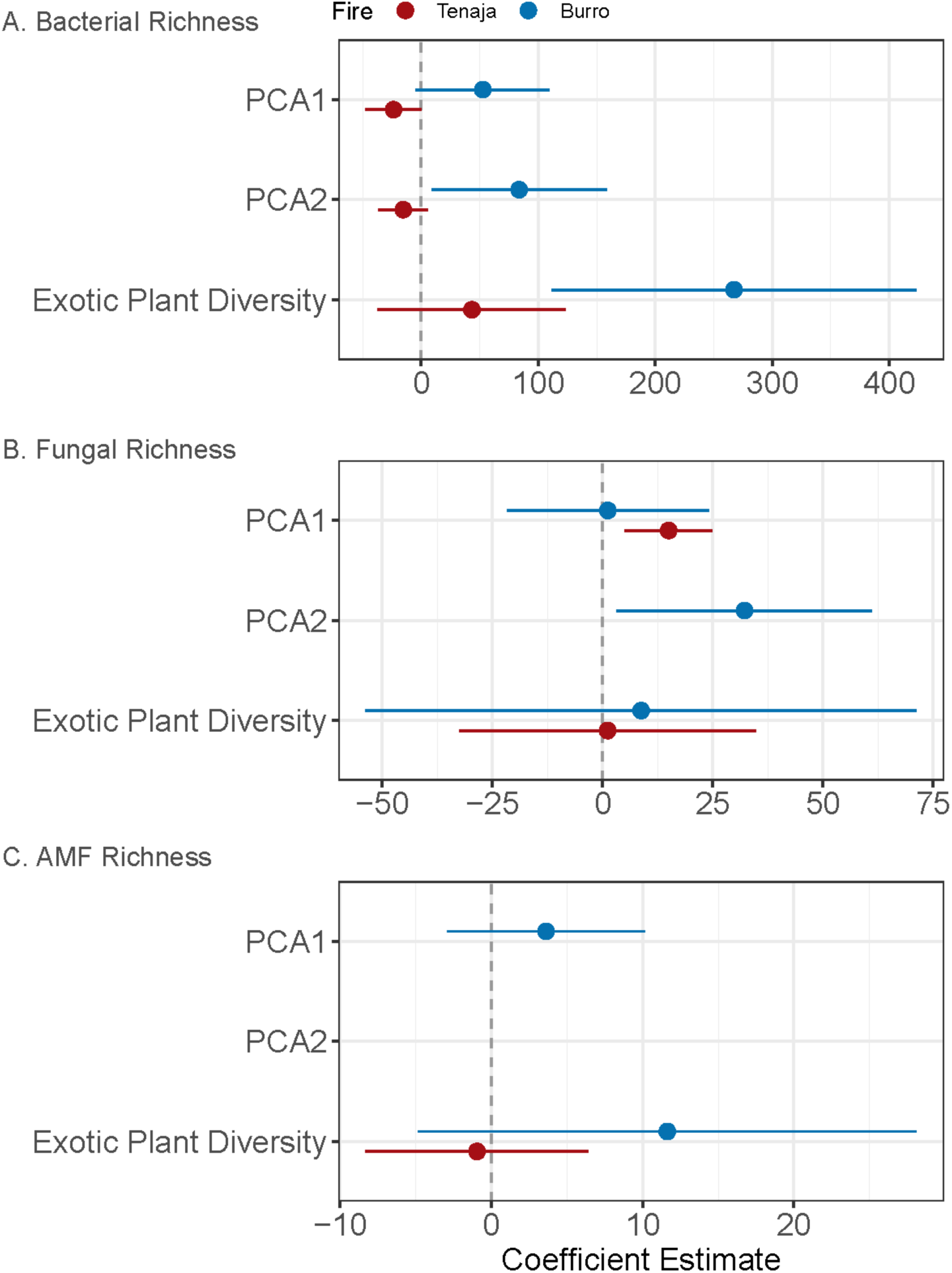
Best fit model parameter coefficients for relating soil variables (PCA1, PCA2) and plant community diversity (exotic Shannon diversity) to richness of A) bacteria B) fungi and C) arbuscular mycorrhizal fungi (AMF) for the Tenaja wildfire versus the Burro prescribed fire. Error bars not overlapping 0 indicate significant parameters in model. Note: the best fit models often contained parameters that were not significant.

**Figure 7.**
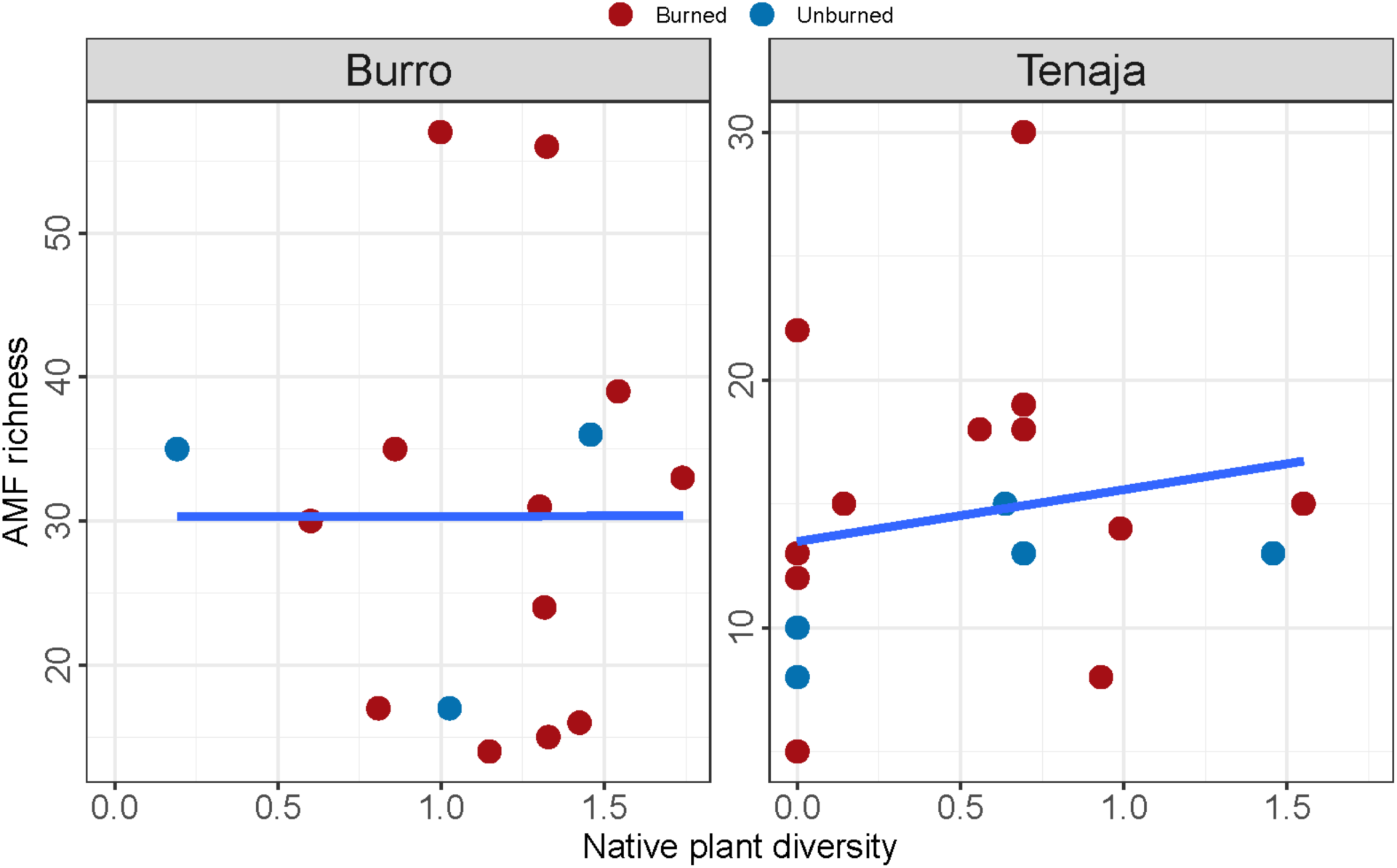
Native plant diversity increases with AMF richness in Tenaja but not Burro fire.

## Discussion

Wildfire had stronger impacts than prescribed fire on microbial communities and both fires had similar impacts on plants. Both fire types significantly reduced the amount of exotic plant cover but neither reduced exotic plant richness (Q1). Regardless of fire type, having a higher cover and diversity of native plants prior to the fire always resulted in higher cover and diversity of native plants post-fire. Soil microbial richness was differentially impacted by fire. Specifically, fungal richness decreased with increasing char levels only within the wildfire but bacterial richness was not impacted by either fire. Bacterial and fungal biomass were impacted by char levels within both the wildfire and prescribed burn (Q2). Both prescribed fire and wildfire significantly altered bacterial and fungal composition, but fungal composition was more altered by wildfire than prescribed fire (Q3). The factors that predicted bacterial, fungal, and AMF richness at the peak of the growing season differed between the two fires (Q4). In the prescribed burn, bacterial and fungal richness was predicted by soil cation exchange capacity and organic matter (PCA2), and bacterial richness was additionally predicted by exotic plant diversity. In the wildfire, bacterial and fungal richness was not related to the plant community but were predicted by soil pH and phosphorus concentrations (PCA1). AMF richness was not related to soil attributes or the exotic plant community but was related to native plant diversity.

Prescribed fire is often used as a tool to reduce exotic abundance and recover native plant species in grasslands (Menke 1992; Dyer 2002; DiTomaso *et al*. 2006). Here, both fires reduced exotic plant cover but neither recovered native plant cover or richness, recapitulating the findings of a recent meta-analysis that neither prescribed burns nor wildfires resulted in increased native plant performance (Alba *et al*. 2015). Appropriately timed fires may reduce viable seeds of exotic plants and allow establishment of native species from the seedbank (Meyer & Schiffman 1999), which not exist. In the prescribed fire site, native forbs like *Deinandra fasciculata* slightly increased in cover in the burned plots. Overall, the prescribed burn site, which has a long history of management, did possess a higher richness and diversity of native species suggesting that periodic fire may be maintaining the integrity of the native seedbank. Yet, in both cases, exotic plant species were more abundant than native, suggesting that regular burning may not necessarily alter the abundance of exotics but may ameliorate the long-term impacts of invasion. For example, long-term plant invasions can erode native seedbanks, reducing the natural colonization of native plant species after management interventions (Cox & Allen 2008). However, a more robust study that directly evaluates seedbank composition across a history of management would provide more insights into this process.

Fires generally have a larger negative impact on fungal than bacterial richness (Pressler, Moore & Cotrufo 2019). Although we know of no similar studies in grasslands, a study in Chinese pine forests found that bacterial richness was not altered by fires of low-, medium-, or high-severity (Li *et al*. 2019). We similarly found that bacterial richness was not impacted by either fire, which is likely due to the high resilience of bacteria that has been detected in forested ecosystems (Xiang *et al*. 2014) and of ammonia oxidizing bacteria in grasslands (Docherty *et al*. 2012). In our study, fungal richness tended to increase in the prescribed fire sites but decreased by 13-17% in the wildfire site. A study of fungal community composition along a boreal coniferous forest fire chronosequence ranging from 2 to 152 years found that fungal richness and diversity was higher at the most recently burned forest sites, whereas the 152 year old site had low diversity and evenness (Sun *et al*. 2015). Thus, it is possible that prescribed fires that are less severe can increase heterogeneity and thereby increase fungal richness as occurs with plants (Agee 1998).

Our findings that wildfire had stronger impacts on soil microbial richness and abundance than prescribed fire and that fire had stronger impacts on fungi than bacteria are largely in line with the literature (Dooley & Treseder 2012; Pressler, Moore & Cotrufo 2019), but there are some key differences. Fire impacts on soil communities are the result of a variety of factors including soil temperature and the inputs associated with post-fire residues such as ash and char (Neary *et al*. 1999). Grassland fires typically are lower severity due to low fuels and quick moving fires and are thus thought to have small impacts on the soil microbial community (Dooley & Treseder 2012). For example, a study examining effects of prescribed fires of varying fire intensity on Northern Great Plain grasslands found no effect of fire on microbial biomass, as measured by PLFA (Reinhart, Dangi & Vermeire 2016), and a meta-analysis of fire effects on biomass found that microbial biomass declined after fires in boreal and temperate forests but not after grassland fires (Dooley & Treseder 2012). Yet, in this case, we were able to detect significant reductions of bacterial and fungal biomass following grassland wildfires. Within both fires fungal and bacterial biomass were sensitive to the amount of char present immediately after the fire. The ability of our study to detect reductions in microbial biomass could be due to our rapid post-fire sampling, usage of char percentage as an index of soil burn severity, which is not typically measured, or because we used an estimate of microbial biomass with higher sensitivity.

The prescribed burn resulted in greater heterogeneity in soil microbial community responses. We observed increased dispersion in fungal composition following the prescribed fire in comparison to the wildfire, which is likely due to the greater variation in remaining litter and char levels in the prescribed fire than the wildfire. Due to our time-series sampling, we observed that the impact of fire on bacterial and fungal abundance tended to fluctuate more over the growing season in the prescribed fire than wildfire site. This may be due to the higher diversity of native plants in the prescribed fire sites or it may be due to the increased heterogeneity after the prescribed fire, since wildfires tend to be more homogenizing (Turner *et al*. 1994).

Both the prescribed fire and wildfire had small but significant impacts on both bacterial and fungal composition. Wildfire had a slightly larger impact on fungal composition than prescribed fire whereas the prescribed fire and wildfire similarly affected bacterial composition, which is likely due to the higher resilience of bacteria to fires (Ferrenberg *et al*. 2013; Xiang *et al*. 2014). There is limited literature for comparison since only 12% of post-fire microbiome studies occur in grasslands. Most studies examine biomass alone (Pressler, Moore & Cotrufo 2019) or arbuscular mycorrhizal colonization and spore counts and not the total fungal community (Cairney & Bastias 2007). However, a recent study investigating the impacts of experimental fire in a California grassland found that fire altered bacterial composition but not alpha diversity (Yang *et al*. 2019), similar to our study. Another recent study of prescribed fires in Mediterranean Aleppo pine forests found that higher severity prescribed fires had larger impacts on bacterial composition than lower severity ones, although both significantly altered bacterial composition (Lucas-Borja *et al*. 2019). Thus, prescribed fires are more likely to alter microbial composition than richness, which could nonetheless alter plant growth and composition (van der Heijden *et al*. 1998).

Primary plant colonizers can dictate the initial microbial community that will subsequently mediate plant recovery (Kardol, Bezemer & van der Putten 2006; Cheeke *et al*. 2019). In the wildfire site with lower overall plant diversity, microbial communities were not sensitive to the aboveground exotic plant community. While we were unable to track the complete recovery of native plant communities in this study, we did find that at the peak of the first post-fire growing season, native plant diversity was positively correlated to AMF richness in the wildfire site. The fact that this positive relationship between AMF richness and native plant richness was not detected in the more biodiverse prescribed fire is corroborated by the fact that the benefit of AMF richness on plant richness tends to saturate after 12-14 AMF species (van der Heijden *et al*. 1998).

Some of the prevalent exotic species present at this site (*Avena* and *Bromus* spp) have exerted strong priority effects on microbial composition in other grasslands within the state (Hausmann & Hawkes 2009; Hausmann & Hawkes 2010), suggesting that relationships between the microbial community and exotic species may become more apparent over time. The low plant diversity of invaded annual grasslands paired with their high productive fine root systems that increase soil organic matter may contribute to lower variation in soil communities (Steenwerth *et al*. 2002), which we observed in the fungal communities of the unburned plots at both sites. Within the prescribed burn that has a history of regularly timed burns, we found that bacterial and fungal richness was sensitive to increasing organic matter, in addition to CEC, and also to soil pH, which is supported by the literature (Fierer & Jackson 2006; Glassman, Wang & Bruns 2017). This site also had greater native plant richness and diversity, potentially increasing the resistance of the AMF community to fire by buffering some of the degradation of mutualistic fungi that can occur with plant invasion (Callaway *et al*. 2008). These findings further suggest that soil legacies associated with long-term invasions may alter the responsiveness of soil microbial communities to the aboveground plant communities.

Unexpected wildfires may provide opportunistic management windows to re-establish native species in degraded grasslands; however, the success of these efforts are contingent on mediating some of the recovery constraints associated with long-term invasion (Larios & Suding 2013). While our findings support the idea that consistent management actions such as periodic prescribed burns may promote native plant diversity (Steenwerth *et al*. 2002), they do not indicate that periodic burns or wildfires will increase native plant performance or cover without intervention. Soil inoculations are likely required to recover native soil microbial and plant diversity in these situations (Aprahamian *et al*. 2016; Emam 2016; Balshor *et al*. 2017), and these strategies are more likely to be successful if local soil inocula are used (Maltz & Treseder 2015) since mycorrhizal partnerships tend to be context dependent (Klironomos 2003). Thus, the success of an opportunistic management action capitalizing on a wildfire would be contingent on addressing invader legacies (e.g. adding seeds, soil amendments) and implementing periodic management actions (e.g. prescribed fires) to regularly clear out persistent plant invaders.

## Acknowledgements

We thank Hailey Laskey, Carole Bell, and Zachary Principe for allowing us access to the Santa Rosa Ecological Plateau Reserve, and Judy Chung, Lachland Charles, Miguel Solis and Soren Weber for aid in field work and plant data collection.

## Authors’ Contributions

SIG conceived of the field experiment with joint project development and experimental design by SIG and LL. JC, JWJR, LL, SIG, SSS conducted field work. JWJR performed molecular work and JC and KEW assisted in bacterial biomass measurements. JC, JWJR, MFPC, and SIG conducted bioinformatics. JWJR, LL, and SIG conducted statistical analyses and made the figures. SIG and LL wrote the first draft. All authors contributed edits to the manuscript and approved the final submission.

## Supplementary Information

**Table S1:** GPS Coordinates and elevation for the 15 sampling locations established at the initial time point for Burro (May 21, 2018) and the 17 sampling locations of Tenaja (October 29th, 2019)

| Site | Line | Plot | Date | Elevation (m) | Latitude | Longitude |
| --- | --- | --- | --- | --- | --- | --- |
| Tenaja | 1 | A | 10/29/19 | 559 | 33.540061 | -117.25194 |
| Tenaja | 1 | B | 10/29/19 | 560 | 33.540235 | -117.252 |
| Tenaja | 1 | C | 10/29/19 | 561 | 33.54031 | -117.25203 |
| Tenaja | 1 | D | 10/29/19 | 563 | 33.541077 | -117.25224 |
| Tenaja | 1 | E | 10/29/19 | 567 | 33.541936 | -117.25248 |
| Tenaja | 2 | A | 10/29/19 | 565 | 33.536947 | -117.25255 |
| Tenaja | 2 | B | 10/29/19 | 566 | 33.537064 | -117.25277 |
| Tenaja | 2 | C | 10/29/19 | 563 | 33.537103 | -117.2528 |
| Tenaja | 2 | D | 10/29/19 | 553 | 33.537698 | -117.25345 |
| Tenaja | 2 | E | 10/29/19 | 547 | 33.538304 | -117.25416 |
| Tenaja | 3 | A | 10/29/19 | 555 | 33.53585 | -117.25527 |
| Tenaja | 3 | B | 10/29/19 | 555 | 33.536004 | -117.25545 |
| Tenaja | 3 | C | 10/29/19 | 555 | 33.536093 | -117.25556 |
| Tenaja | 3 | D | 10/29/19 | 551 | 33.536601 | -117.25618 |
| Tenaja | 3 | E | 10/29/19 | 549 | 33.537199 | -117.25698 |
| Burro | 1 | O | 5/21/18 | 592 | 33.526254 | -117.22979 |
| Burro | 1 | A | 5/21/18 | 592 | 33.526324 | -117.22973 |
| Burro | 1 | B | 5/21/18 | 592 | 33.526215 | -117.22982 |
| Burro | 1 | C | 5/21/18 | 592 | 33.526104 | -117.2299 |
| Burro | 1 | D | 5/21/18 | 593 | 33.525502 | -117.23036 |
| Burro | 1 | E | 5/21/18 | 597 | 33.524726 | -117.23095 |
| Burro | 2 | O | 5/21/18 | 594 | 33.525167 | -117.22533 |
| Burro | 2 | A | 5/21/18 | 594 | 33.525239 | -117.22527 |
| Burro | 2 | B | 5/21/18 | 593 | 33.525143 | -117.22536 |
| Burro | 2 | C | 5/21/18 | 594 | 33.525041 | -117.22547 |
| Burro | 2 | D | 5/21/18 | 595 | 33.524476 | -117.22603 |
| Burro | 2 | E | 5/21/18 | 596 | 33.523786 | -117.22668 |
| Burro | 3 | O | 5/21/18 | 603 | 33.525204 | -117.22766 |
| Burro | 3 | A | 5/21/18 | 602 | 33.52529 | -117.22759 |
| Burro | 3 | B | 5/21/18 | 604 | 33.525163 | -117.22768 |
| Burro | 3 | C | 5/21/18 | 603 | 33.525043 | -117.22775 |
| Burro | 3 | D | 5/21/18 | 604 | 33.524426 | -117.22818 |
| Burro | 3 | E | 5/21/18 | 605 | 33.523635 | -117.22871 |

**Table S2.**
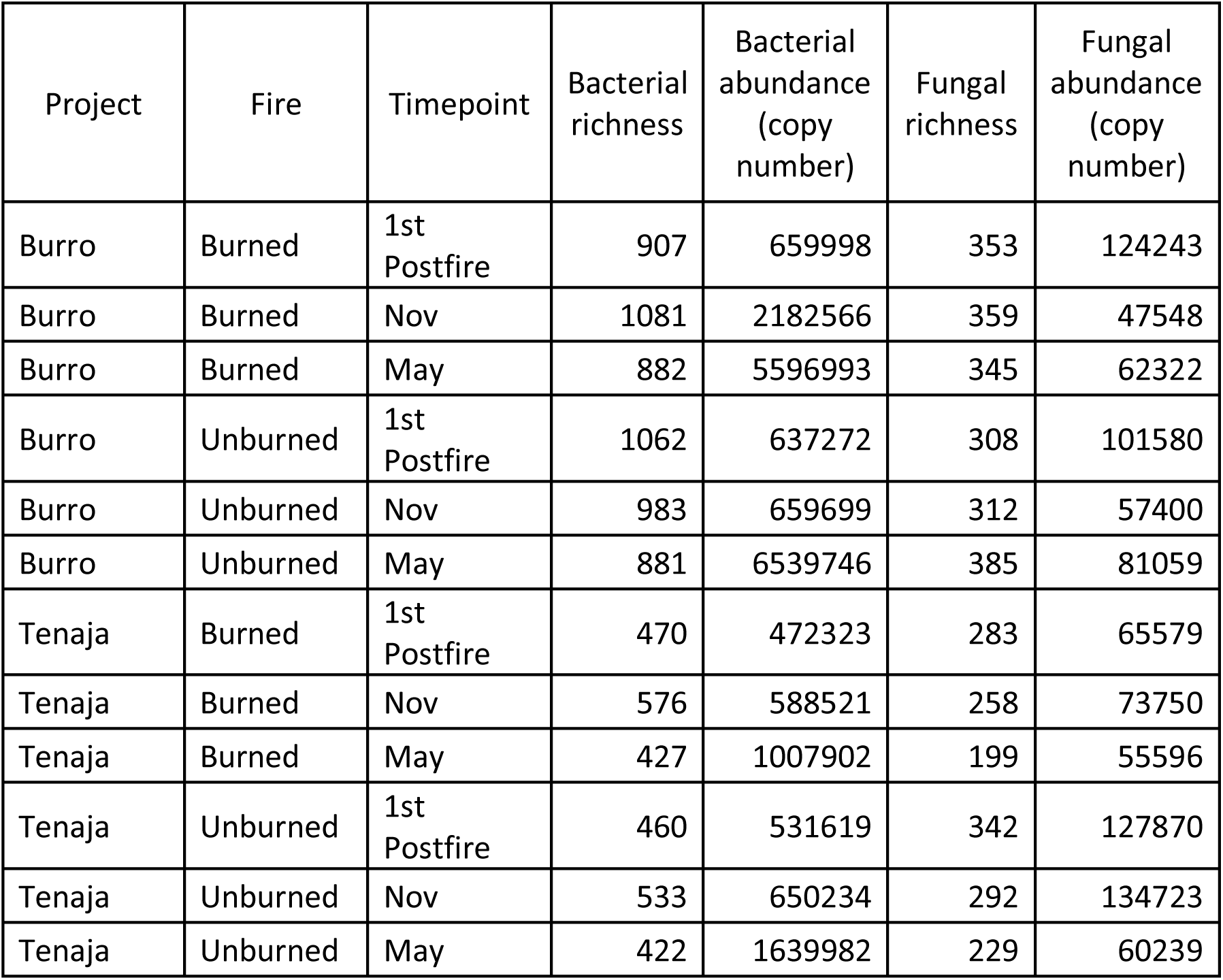
Summary of mean bacterial and fungal richness (observed species number after rarefaction) and abundance (copy number) for the prescribed (Burro) versus wildfire (Tenaja) in burned versus unburned plots in the three post-fire time points.

| Project | Fire | Timepoint | Bacterial richness | Bacterial abundance (copy number) | Fungal richness | Fungal abundance (copy number) |
| --- | --- | --- | --- | --- | --- | --- |
| Burro | Burned | 1st Postfire | 907 | 659998 | 353 | 124243 |
| Burro | Burned | Nov | 1081 | 2182566 | 359 | 47548 |
| Burro | Burned | May | 882 | 5596993 | 345 | 62322 |
| Burro | Unburned | 1st Postfire | 1062 | 637272 | 308 | 101580 |
| Burro | Unburned | Nov | 983 | 659699 | 312 | 57400 |
| Burro | Unburned | May | 881 | 6539746 | 385 | 81059 |
| Tenaja | Burned | 1st Postfire | 470 | 472323 | 283 | 65579 |
| Tenaja | Burned | Nov | 576 | 588521 | 258 | 73750 |
| Tenaja | Burned | May | 427 | 1007902 | 199 | 55596 |
| Tenaja | Unburned | 1st Postfire | 460 | 531619 | 342 | 127870 |
| Tenaja | Unburned | Nov | 533 | 650234 | 292 | 134723 |
| Tenaja | Unburned | May | 422 | 1639982 | 229 | 60239 |

**Table S3:** Model summary statistics for plant community composition metrics for exotic and native species.

| Table 1. Model summary statistics for plant community composition metrics for exotic and native species. |  |  |  |  |  |  |
| --- | --- | --- | --- | --- | --- | --- |
| Diversity Metric | Burn Treatment |  | Fire Site |  | Fire*Burn |  |
|  | F | p- value | F | p- value | F | p- value |
| Exotic Cover | 4.15<sub>1,28</sub> | 0.05 | 0.60<sub>1,28</sub> | 0.44 | 0.52<sub>1,28</sub> | 0.47 |
| Exotic Diversity | 1.04<sub>1,15</sub> | 0.32 | 17.05<sub>1,16.6</sub> | <0.001 | 0.39<sub>1,16.6</sub> | 0.54 |
| Exotic Richness | 0.12<sub>1,27.3</sub> | 0.73 | 7.79<sub>1,27.3</sub> | 0.009 | 0.09<sub>1,27.3</sub> | 0.77 |
| Native Cover | 0.26<sub>1,15</sub> | 0.62 | 33.7<sub>1,16</sub> | <0.001 | 0.90<sub>1,16</sub> | 0.36 |
| Native Diversity | 0.61<sub>1,27.3</sub> | 0.44 | 7.92<sub>1,27.3</sub> | <0.01 | 0.82<sub>1,27.3</sub> | 0.37 |
| Native Richness | 0.24<sub>1,15</sub> | 0.63 | 12.34<sub>1,16</sub> | <0.01 | 0.19<sub>1,16</sub> | 0.67 |

**Table S4:** Model summary statistics for bacterial and fungal richness and abundance for Burro and Tenaja fires. Predictor variables included Initial char cover, time, and their interaction. Plot number was included as a random effect, with an autoregressive correlation structure.

|  | Bacterial |  |  |  | Fungal |  |  |  |
| --- | --- | --- | --- | --- | --- | --- | --- | --- |
|  | Richness |  | Abundance |  | Richness |  | Abundance |  |
|  | F | p-value | F | p-value | F | p-value | F | p-value |
| **Burro** |  |  |  |  |  |  |  |  |
| Initial Char | 0.39<sub>1,1</sub><br>3 | 0.55 | 0.14<sub>1,13</sub> | 0.71 | 0.15<sub>1,13</sub> | 0.70 | 1.49<sub>1,13</sub> | 0.24 |
| Time | 3.50<sub>2,2</sub><br>6 | <0.05 | 1.71<sub>2,26</sub> | 0.20 | 0.11<sub>2,26</sub> | 0.89 | 29.90<sub>2,2</sub><br>6 | <0.001 |
| Char*Time | 1.91<sub>2,2</sub><br>6 | 0.17 | 2.86<sub>2,26</sub> | 0.08 | 0.97<sub>2,26</sub> | 0.39 | 5.88<sub>2,26</sub> | <0.01 |
| **Tenaja** |  |  |  |  |  |  |  |  |
| Initial Char | 0.17<sub>1,2</sub><br>9 | 0.68 | 7.78<sub>1,29</sub> | <0.01 | 22.41<sub>1,2</sub><br>9 | <0.001 | 6.33<sub>1,29</sub> | <0.05 |
| Time | 7.32<sub>2,2</sub><br>9 | <0.01 | 29.12<sub>2,2</sub><br>9 | <0.001 | 16.41<sub>2,2</sub><br>9 | <0.001 | 6.04<sub>2,29</sub> | <0.01 |
| Char*Time | 0.18<sub>2,2</sub><br>9 | 0.83 | 1.40<sub>2,29</sub> | 0.26 | 0.57<sub>2,29</sub> | 0.57 | 0.96<sub>2,29</sub> | 0.40 |

**Figure S1.**
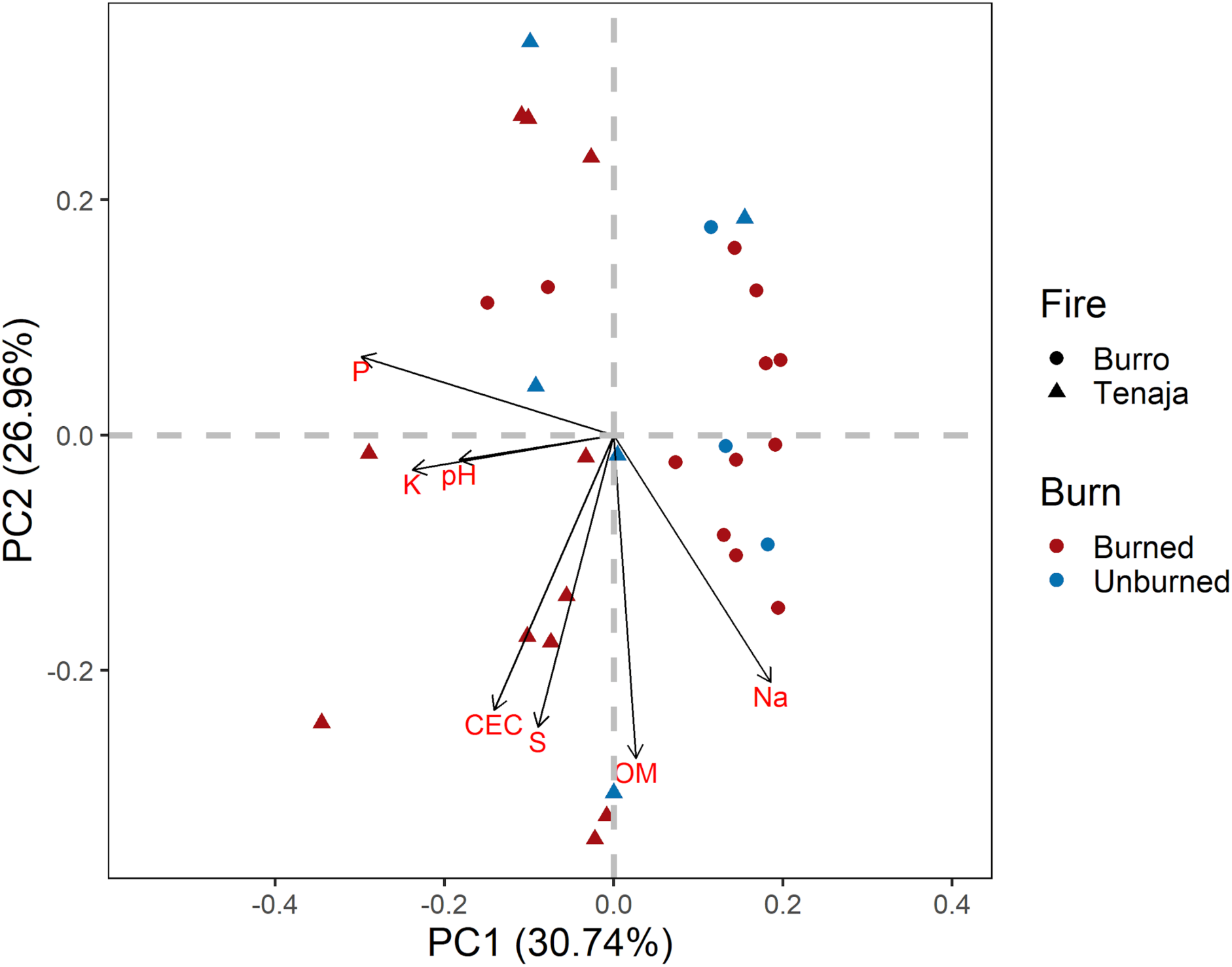
Principal component analysis of soil abiotic variables used to describe soil environmental gradient used for analyses. For PC1, increasing values indicate greater sodium (Na) concentrations and lower values indicate higher soil pH, phosphorus (P) and Potassium (K). For PC2 increasing values indicate lower cation exchange capacity (CEC) while negative values indicated plots with high CEC, organic matter (OM) and sulfur (S).

**Fig. S2.**
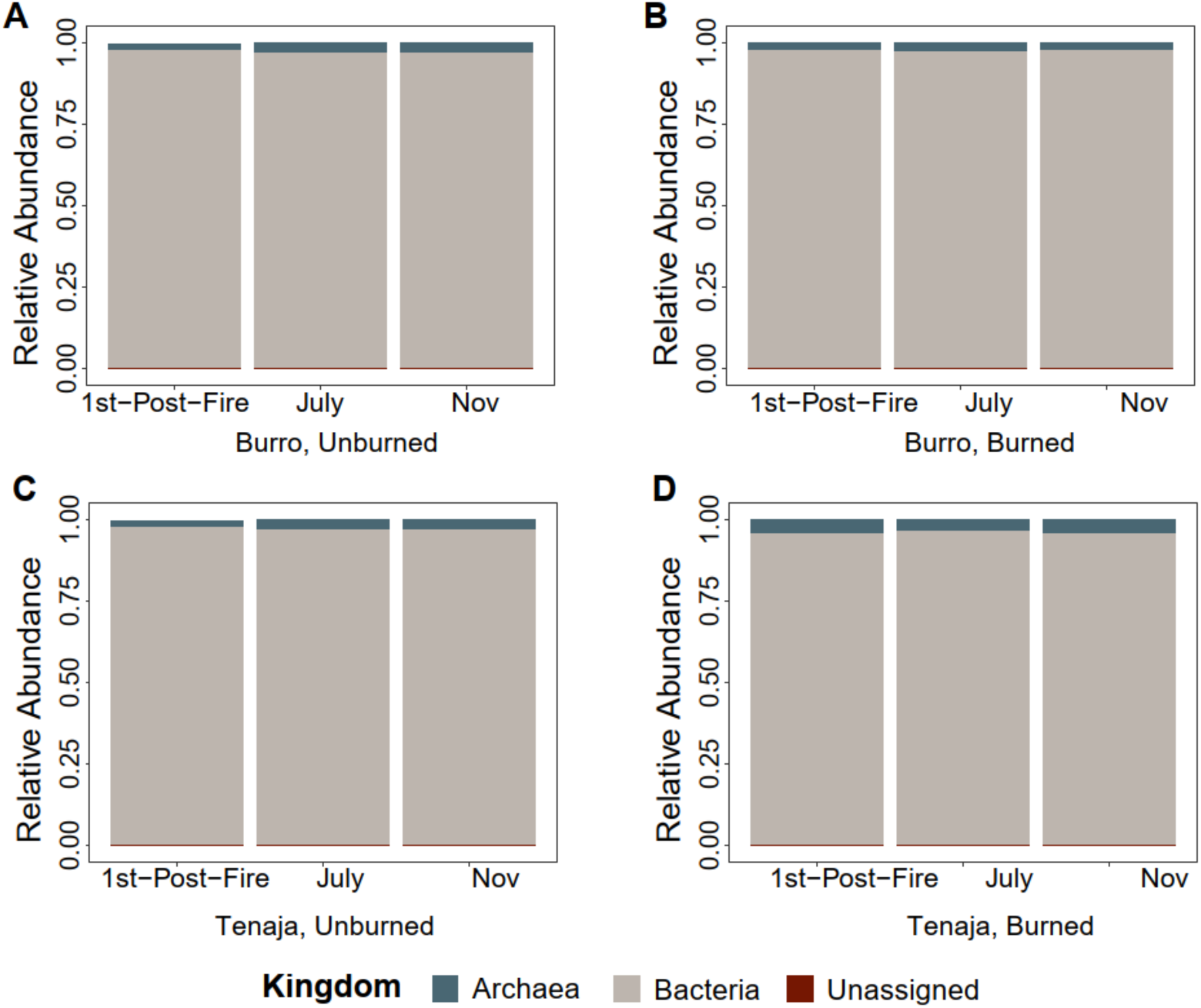
Relative sequence abundance of archaeal versus bacterial reads in the Burro prescribed fire A) unburned plots and B) burned plots and in the Tenaja wildfire C) unburned and D) burned plots at the three time points.

**Fig. S3.**
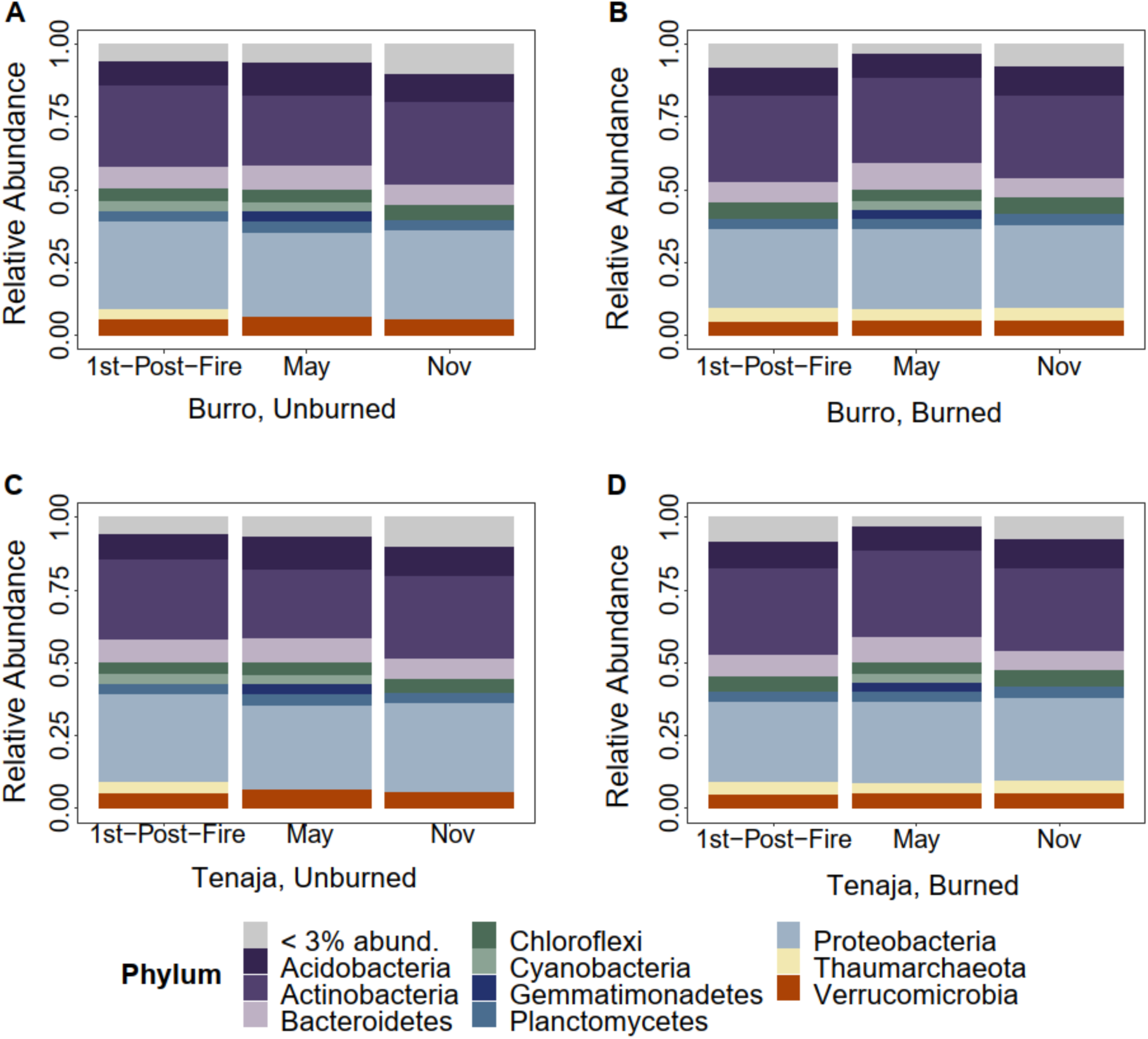
Relative sequence abundance of bacterial phyla in the Burro prescribed fire A) unburned plots and B) burned plots and in the Tenaja wildfire C) unburned and D) burned plots at the three time points.

**Fig S4.**
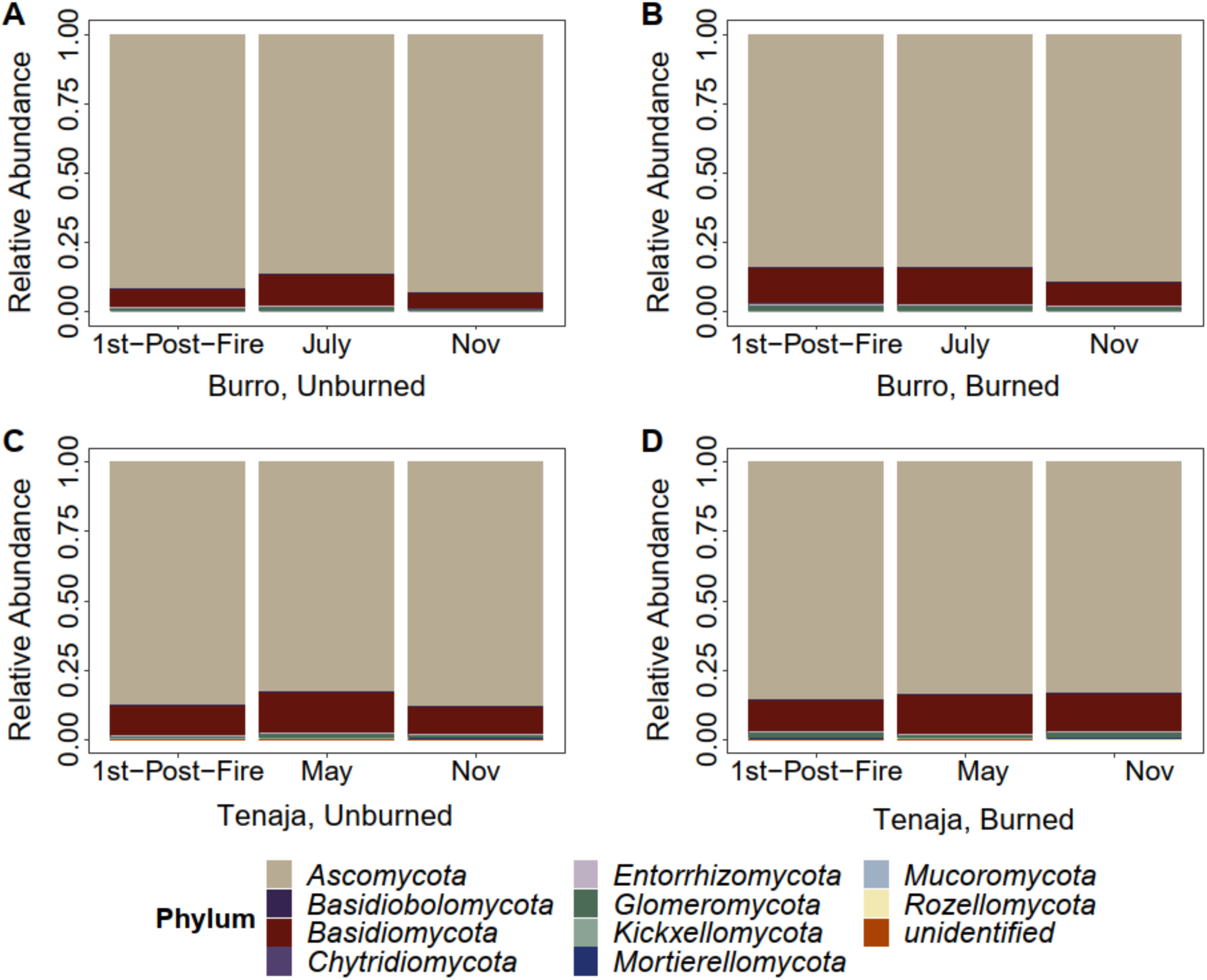
Relative sequence abundance of fungal phyla in the Burro prescribed fire A) unburned plots and B) burned plots and in the Tenaja wildfire C) unburned and D) burned plots at the three time points.

**Fig S5.**
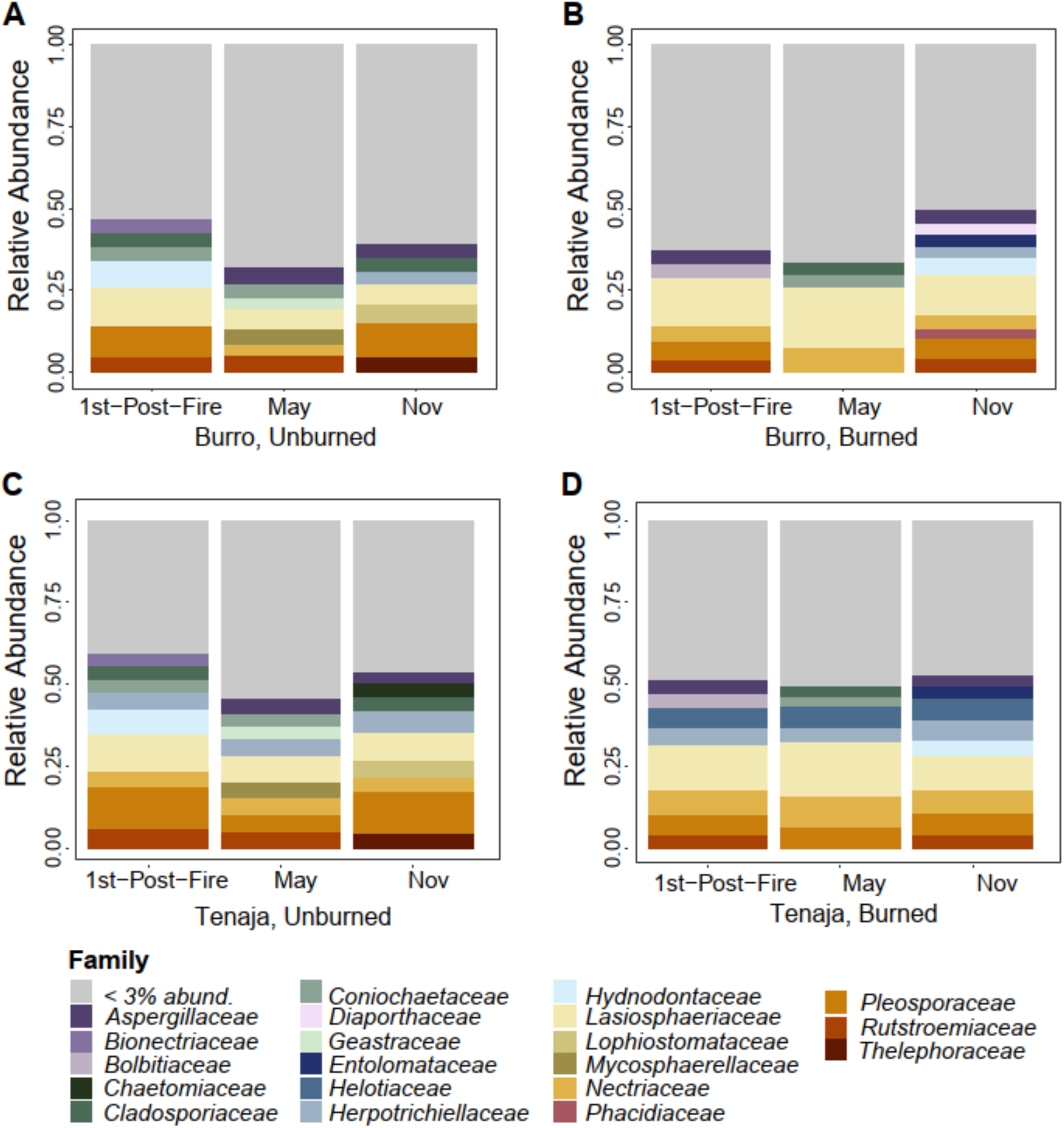
Relative sequence abundance of fungal families in the Burro prescribed fire A) unburned plots and B) burned plots and in the Tenaja wildfire C) unburned and D) burned plots at the three time points. Burro samples and in C) unburned Tenaja versus D) burned Tenaja samples.

**Methods S1:** Description of molecular methods for obtaining 16S and ITS amplicon sequencing libraries.

For fungi, we combined the gene-specific primers (ITS4-fun and 5.8s; Taylor *et al*. 2016) at 0.5 μl each at 10µM, 5 µl of undiluted fungal DNA, 6.5 µl of Ultra-Pure Sterile Molecular Biology Grade (Genesee Scientific, San Diego, CA, USA) water and 12.5 µl of AccuStart ToughMix (Quantabio, Beverly, MA, USA). Thermocycler conditions were: 94 °C for 2 min, followed by 31 cycles of 94 °C for 30 s, 55 °C for 30 s, 68 °C for 2 min followed by a 10 min extension at 68 °C. For bacteria, we combined 1 μl of 1:10 diluted DNA, 10.5 μl of water, 12.5 μl of AccuStart ToughMix and 0.5 μl each of the 10 uM 515F-806R primers. Thermocycler conditions were: 94°C for 2 min, followed by 30 cycles of 94°C for 30 s, 55°C for 30 s, 68°C for 1 min followed by a 2 min extension step for at 68°C. Both conditions ended with a 4°C hold. PCR products were then cleaned with AMPure XP magnetic Bead protocol (Beckman Coulter Inc, Brea, CA, USA) following manufacturers protocols. The DIP PCR2 primers containing the barcodes and adaptors for Illumina sequencing were ligated to the amplicons during the second PCR step in a 25 µL reaction containing 2.5 μl of the 10 uM DIP PCR2 primers, 6.5 μl of water, 12.5 μl of Accustart ToughMix and 1 μl of PCR 1 product. Thermocycler conditions for the second PCR were: 94°C for 2 min followed by 10 cycles of 94°C for 30 s, 60°C for 30 s, 72°C for 1 min, and finally ending at a 4°C hold. Bacterial and fungal PCR products were then separately pooled based on gel electrophoresis band strength and cleaned with AMPURE following established methods (Glassman *et al*. 2018). The 16S and ITS pools were each checked for quality and quantity with an Agilent 2100 Bioanalyzer, then pooled at 0.4 bacteria to 0.6 fungi ratio prior to sequencing.

**Methods S2**. Methodological details for qPCR reactions to obtain estimates of bacterial and fungal biomass

Bacterial biomass was estimated based on bacterial 16S rRNA genes using the Eub338/Eub518 primer set (Fierer *et al*. 2005) and fungal biomass was estimated based on fungal 18S rRNA genes using the FungiQuant-F and FungiQuant-R primer set (Liu *et al*. 2012). qPCR reactions were performed in triplicate with 1 µL of undiluted DNA added to 9 µl of qPCR master mixer containing 1 µl of 0.05M Tris-HCl ph8.3, 1 µL of 2.5mM MgCl_2_ (New England BioLabs; NEB; Ipswich, MA, USA), 0.5 µL of 0.5mg/ml BSA, 0.5 µL of 0.25mM dNTPs (NEB), 0.4 µL of both forward and reverse primer at 0.4µM, 0.5 µL of 20X Evagreen Dye (VWR), 0.1 µL of Taq DNA polymerase (NEB) and the remaining volume of 4.6 µL with the molecular grade water. Each reaction was run in triplicate in 384 well plates on CFX384 Touch Real-Time PCR Detection System starting at 94°C 5 min, followed by 40 cycles of a denaturing step at 94°C 20 sec, primer annealing at 52°C for bacteria or 50°C for fungi at 30 sec, and an extension step at 72°C for 30 sec. Standards were generated by cloning the 18S region of *Saccharomyces cerevisiae* or the 16S region of *Escherichia coli* into puc57 plasmid vectors, which were constructed by GENEWIZ, Inc. (NJ, USA) as previously established (Averill & Hawkes 2016) Melt curves were generated and copy number extracted using the following equation: 10^(Cq-b)/m^ where Cq is the average of 3 technical replicates of Cq value and b is the y intercept and m is the slope.

